# CtBP2 triggers CCN1-induced metastatic dissemination of osteosarcoma cells in a non-hypoxic microenvironment

**DOI:** 10.1101/2023.12.07.570584

**Authors:** Laura Di Patria, Nadia Habel, Robert Olaso, Bojana Stefanovska, Olivia Fromigue

**Author notes:** Corresponding Author : Dr Olivia Fromigué, Inserm UMR981, Gustave Roussy Cancer Campus, 39 Rue Camille Desmoulins, F-94805 Villejuif, France. Centre de Traitement de l’Information Génétique (CTIG), INRAE, Jouy en Josas, France. Department of Biochemistry and Structural Biology, Howard Hughes Medical Institute, University of Texas Health San Antonio, San Antonio, TX, USA.

## Abstract

Osteosarcoma is the most prevalent pediatric solid bone tumor. These tumors are highly metastatic and frequently develop resistance to chemotherapy, leading to poor survival rate for patients. We found that C-terminal Binding Protein 2 (CtBP2) and Cysteine-rich protein 61 (CYR61/CCN1) expression level correlated positively in a panel of cell lines. *In silico* analysis of protein-protein interaction network revealed a link with stemness markers. We confirmed that CtBP2 influences stemness markers expression, cell clonogenicity, cell migration, matrix metalloproteinase activity and cell invasion. Surprisingly, using syngeneic tumor cells graft models, while induction of CtBP2 expression correlated with the metastatic dissemination process, it occurred only at the invasive front. Hypoxic conditions in central tumor region interfered with CtBP2 induction. Globally, we identify for the first time that CtBP2 is a required inducing factor in the CYR61-related metastatic progression of osteosarcoma. Moreover, we demonstrate that while CtBP2 is a downstream transcriptional target of CYR61 signaling cascade, it occurs only under non-hypoxic conditions. The present study suggests that CtBP2 may represent a potential pivotal target for therapeutic management of metastases spreading in osteosarcoma.

## 1. Introduction

Osteosarcoma is the most common malignant skeletal tumor and most often affects children and young adults. These bone tumors frequently spread locally, and develop metastases into distant organs (predominately to the lungs)(*1*). The therapeutic protocol consists of neo-adjuvant chemotherapy, surgical removal of the tumor if the location is accessible, then multi-agent chemotherapy regimens. The 5-year event free survival (EFS) and overall survival (OS) rates for patients with localized osteosarcoma are 64% and 90%, respectively (*2*). In contrast, patients with detectable metastases at diagnosis or with recurrent disease have worse overall prognosis with 5-year EFS and OS rates of only 25% and 21%, respectively. The improvement of clinical outcome for patients with poor prognosis is challenging, and requires the establishment of new therapeutic strategies to prevent the development of metastatic disease.

Several key processes are required for the successful establishment of distant metastasis. Thereby, early in the metastatic cascade, cancer cells must acquire invasive properties, and gain access to the blood or lymphatic vascular systems. We showed previously that statins induce anti-tumor effects on osteosarcoma cells (*3–5*), and lead to the down-regulation of cysteine rich protein 61 (CYR61/CCN1) expression levels in these tumor cells (*6*). The immediate-early gene *cyr61* encodes a member of the extracellular matrix-associated CCN family of six homologous cysteine-rich proteins in vertebrates comprising connective tissue growth factor (CTGF), nephroblastoma overexpressed (NOV), and Wnt-induced secreted proteins (WISPs). This family of proteins, with CYR61 as forerunner, are involved in multiple physiological functions such as skeletal and cardiovascular development and injury repair (*7–10*). In different solid cancer types, CYR61 promotes tumor growth and vascularization as well as cell invasiveness and metastasis (*11–15*). We established that CYR61 expression levels correlate with the metastatic capacities of osteosarcoma tumor cells both *in vitro* and in preclinical murine models, and correlate positively with tumor grade in patients (*6*). Our previous data support the main role of CYR61 in tumor vascularization through the promotion of a neo-angiogenesis and remodeling/destruction of the extracellular matrix surrounding the primary tumor (*6, 16*). In addition, we demonstrated that CYR61 silencing significantly reduces *in vitro* cell migration potential and expression of pro-angiogenic factors, but also reduces *in vivo* tumor neovascularization and lung metastases occurrence (*6, 16, 17*). The overexpression of CYR61 in cancer cells is commonly associated with epithelial-to-mesenchymal transition (EMT). In fact, when progressing to a more malignant state, tumor cells frequently lose some of their epithelial characteristics and acquire properties of migrating cells (*18*). We have reported a process similar to EMT in osteosarcoma cells despite their mesenchymal origin, and demonstrated that CYR61 silencing reinforces the mesenchymal phenotype (*17*). Globally, our data established CYR61 as promising key marker of tumor dissemination thus representing an appealing therapeutic target in metastatic osteosarcoma. However, its ubiquitous nature and its multiple physiological roles are the main obstacles for the development of targeted therapeutic strategies. A better characterization of the pro-metastatic cascade reliant on CYR61 is crucial to refine the therapeutic options.

In the present study, we substantiated the effect of CYR61 on dissemination abilities of osteosarcoma cells, and identified a new essential downstream effector. Our results indicate that the expression of the transcriptional co-repressor CtBP2 (Carboxyl-terminal Binding Protein 2) is induced by CYR61 at a transcriptional level. Furthermore, we highlighted a spatial regulation of CtBP2 induction, mainly dependent on oxygen levels. Our results demonstrate that CtBP2 is an essential component of the CYR61-dependent pro-metastatic cascade in osteosarcoma cells.

## 2. Materials and Methods

### 2.1. Maintenance of osteosarcoma cell lines

The murine K7M2 cell line, and the human 143B, HOS, MG-63, SaOS-2, and U-2 OS cell lines were originally obtained from ATCC (LGC Standards sarl, Molsheim, France). The human IOR/OS18 cell line was kindly provided by Pr M Serra (Istituti Ortopedici Rizzoli, Bologna; Italy) (*19*). The human CAL72 cell line was kindly provided by Dr N Rochet (Institute of Biology Valrose, Nice, France) (*20*). The human OHS-4 cell line was kindly provided by Drs Fournier and Price (University of California, San Diego, CA, USA) (*21*).

All cell lines were routinely grown in GlutaMAX^TM^-containing high glucose Dulbecco’s Modified Eagles Medium (DMEM; Thermo Fisher Scientific, Courtabœuf, France) supplemented with 10% heat inactivated fetal bovine serum (FBS), at 37°C in a saturated humidity atmosphere (5% CO_2_ and 95% air). Culture media were changed three times a week, and regularly tested for mycoplasma contamination (PCR-based assay from Minerva-Biolabs, Berlin, Germany).

### 2.2. Generation of stable cell lines by lentiviral vector transduction

Ready-to-use lentiviral particles containing a vector encoding human or murine CtBP2 shRNA sequences or non-targeting (control) shRNA sequences were purchased from Santa Cruz Biotechnology (Santa Cruz, CA, USA). Briefly, sub-confluent target cells were incubated with particles and 4 µg/mL polybrene (Hexadimethrine bromide; Sigma-Aldrich, Lyon, France) for 18 h, washed once with medium, and cultured for 48 h. Non-transduced cells were eliminated by puromycin dihydrochloride selection (10 µg/mL; Sigma-Aldrich) for 3 days. Viable cells were routinely maintained in complete medium.

Lentiviral vectors for CtBP2 over-expression in mammalian cells were generated by Gateway cloning. The donor vector pDONR223-CtBP2 was purchased from the central repository DNASU (Arizona State University, Tempe, AZ, USA), and the destination lentiviral empty vector pLX303 was purchased from Addgene (Cambridge, MA, USA). The sub-cloning of the CtBP2 sequence into pLX303 lentiviral vector was performed by site-specific recombination using LR clonase-II (Thermo Fisher Scientific), according to the manufacturer’s recommendations. Particles were produced using human embryonic kidney cells HEK293T as previously described (*22*). Supernatants were collected after 48 h incubation, centrifuged at 200 x *g* to remove cell debris, and filtered through a 0.45 µm low protein-binding filter (Corning, Bath, UK). Sub-confluent target cells were incubated with the particles-containing supernatant supplemented with 4 µg/mL polybrene for 18 h, washed once with medium, and cultured for 48 h before Blasticidin selection (5 µg/mL; Sigma-Aldrich) for 3 days. Viable cells were routinely maintained in complete medium.

The K7M2 cell lines stably modified for CYR61 were established as previously described (*6*).

### 2.3. Cell cytoplasmic labelling

The cell fluorescent labelling was performed using CytoLabeling reagent dyes (Abcam, Cambridge, UK) according to the manufacturer’s recommendations.

### 2.4. RNA extraction

Total RNAs were isolated using the TRIzol RNA Isolation Reagent (Thermo Fisher Scientific), and stored at -80°C. RNA quantitation, and quality control were performed on Agilent 2100 Bioanalyzer to select samples exhibiting RNA Integrity Number >8.

### 2.5. Reverse Transcription and quantitative Polymerase Chain Reaction (RT-qPCR)

Total RNA (3 µg) were denatured for 10 min at 70°C then reverse transcribed at 37°C for 90 min in a buffer containing Superscript II reverse transcriptase (Thermo Fisher Scientific), DTT (10 mM), random hexanucleotides (2.5 µM), and dNTPs mix (0.5 mM). A final step of inactivation of the enzyme was carried out at 95°C for 5 min.

Real-time quantitative Polymerase Chain Reaction (qPCR) was carried out on ViiA7 apparatus (Thermo Fisher Scientific) using SYBR Green Master kit (Thermo Fisher Scientific) supplemented with 0.5 µM of specific primers (sequences are detailed in supplemental Tables S1 and S2), and thermal conditions: enzyme activation at 95°C for 15 min then 45 cycles of: denaturation at 95°C for 20 sec, annealing at 58°C for 20 sec and extension at 72°C for 20 sec. Upon run completion, a melting curve analysis was included to assure that only one PCR amplicon was formed. The relative fold gene expressions were estimated using the 2^-ΔΔCt^ method.

### 2.6. Cell metabolic activity assay

The metabolic activity was evaluated using a colorimetric assay. Briefly, MTT reagent (3-(4,5-dimethylthiazol-2-yl)-2,5-diphenyltetrazolium bromide; Sigma Aldrich) was added to the culture medium at a final concentration of 1 mg/mL for the last 1 h of incubation. The absorbance of tetrazolium precipitates solubilized in DMSO was measured at the wavelength of 592 nm (Victor-X2 Microplate Reader, PerkinElmer France SAS, Villebon sur Yvette, France).

### 2.7. DNA replication assay

The cell proliferation rate was evaluated using the Kit Biotrak ELISA System (GE Healthcare, Orsay, France) according to the manufacturer’s instructions. Briefly, BrdU reagent (5-bromo-2’-deoxyuridine) was added to the culture medium at a final concentration of 10 µM for the last 3 h of incubation. The quantification of BrdU incorporation into newly synthesized DNA of the actively proliferating cells was measured at the wavelength of 592 nm.

### 2.8. Clonogenic assay

The ability of single cells to grow into colonies was evaluated using an *in vitro* cell survival assay. Cells were seeded at the density of 100 cells/cm² in 12-wells plates, and cultured for 10 days. Cells were then washed with PBS, fixed with 75% ethanol, and stained with crystal violet solution (0.05% in ethanol). The number and size of cell colonies were then evaluated using ImageJ software (v 1.53t; National Institutes of Health, Bethesda, MD, USA).

### 2.9. Wound healing assay

The cell motility was evaluated using removable silicone inserts, according to manufacturer’s instructions (Ibidi, Martinsried, Germany). Cell monolayers were cultured for 16 h, fixed in 75% ethanol, then stained with crystal violet solution (0.05% in ethanol). Recovery of the denuded area was computerized using an inverted digital microscope (EVOS), and evaluated using ImageJ software package. Lesion area surface at time zero was used as matrix.

### 2.10. Chemotactic cell migration and invasion assays

The cell migration rate was evaluated using modified Boyden chamber (8 μm pore size Falcon^®^ Transwell inserts, Corning, Boulogne-Billancourt, France). Briefly, 5×10^4^ serum-starved cells were seeded in the insert, placed in wells filled with 10% FCS containing medium as attractant. After 10 h incubation, the cells remained in the upper chamber were carefully removed using a cotton swab, whereas the cells that have migrated through the membrane are fixed, stained with crystal violet solution and counted.

The cell invasion rate was evaluated using the same protocol except the use of Matrigel® Basement Membrane Matrix-coated modified Boyden chamber.

### 2.11. Matrix Metalloproteinase activity assay

Cells were lysed in 0.1 M Tris/HCl (pH 7.5), 0.1% Tween-80, on ice for 15 min, then centrifuged for 5 min at 10,000 x *g* to discard insoluble materials. MMP2 activity was evaluated by a colorimetric assay using Ac-Pro-Leu-Gly-[2-mercapto-4-methyl-pentanoyl]-Leu-Gly-OC2H5 thiopeptide (50 µM; BioMol International, Hamburg, Germany) in 50 mM Hepes, 10 mM CaCl_2_, 1 mM ZnCl_2_, 0.05% Brij35, 1 mM DTNB buffer according to the manufacturer’s recommendations. The activity was expressed as treated over control ratio after correction for total protein content.

### 2.12. Multicellular Tumor Spheroids

Three-dimensional (3D) culture invasion was assessed using small cell aggregates. Cells were seeded at 5×10^3^ cells/50 µL/well of 96-well round-bottomed plates coated with Poly-2-hydroxyethyl methacrylate (12mg/ml; Sigma-Aldrich), and centrifuged at 200 x *g* for 5 min to form clusters. The next day, cells were self-assembled in 3D spheroids with average diameter of 188.5 ± 8.2 µm. At day 2, the medium was supplemented with Matrigel® Basement Membrane Matrix (3.6 mg/mL) to allow cell invasion. Immunofluorescence images were acquired on a Leica epifluorescence inverted microscope (Leica Microsystems Ltd., Wetzlar Germany) equipped with a sCMOS camera and a PeCon chamber i8 to perform live imaging at 37°C with 5% CO_2_. The microscope was steered by the Leica dedicated LasX software. Images were captured at different time points using All quantifications were done with ImageJ software.

Larger spheroids were prepared using 5×10^4^ cells/well, cultured for 10 days in complete medium, fixed in 4% paraformaldehyde then embedded in paraffin.

### 2.13. Western blotting

Whole cell lysates were prepared by homogenization in buffer containing 50 mM Tris/HCl pH 7.4; 150 mM NaCl; 10 mM MgCl_2_; 1% Igepal; 10% Glycerol; 2 mM activated Na_3_VO_4_, and cocktail of proteases inhibitors (Sigma Aldrich), then centrifugation at 10,000 x *g* for 15 min to remove insoluble material. Protein concentration was determined using a colorimetric assay based on the Bradford dye-binding method (Bio-Rad Protein Assay Dye Reagent, Bio-Rad Laboratories, Marnes-la-Coquette, France). Aliquots of 30 µg proteins were separated by SDS–PAGE (sodium dodecyl sulfate–polyacrylamide gel electrophoresis), then transferred to nitrocellulose membranes (Whatman-GE Healthcare Life Sciences, Buckinghamshire, UK). The membranes were incubated for 2 h at RT in casein blocking buffer (Sigma Aldrich), then overnight at +4°C with rabbit anti-CtBP2 antibody (Abcam #ab128871; diluted 1:1000), rabbit anti-CYR61 (Bio-Techne SAS #NB100-356; diluted 1:1000) or rabbit anti-beta Actin (Sigma #; diluted 1:2000). After three washes with TBST buffer [50 mM Tris/HCl pH 7.4, 150 mM NaCl, 0.1% (v/v) Tween-20], membranes were incubated with horseradish peroxidase-conjugated secondary antibodies (1:20,000 dilution in blocking buffer) for 1 h at RT. After final washes, the blots were developed using enhanced chemiluminescence western blotting detection reagent (Thermo Fisher Scientific). Signals were captured using CCD camera of the ChemiDoc XRS^+^ system (BioRad Laboratories), and quantified using the ImageJ software.

### 2.14. Syngeneic tumor cells graft models

Animal experiments and procedures were conducted according to the guidelines formulated by the European Commission for experimental animal use (L358-86/609EEC), and with the approval of the local ethical animal committee of Paris Saclay University (CEEA 26, Villejuif, France). *In vivo* assays were performed as previously described (*16*). Briefly, 5-weeks old female BALB/c *nude* mice (Charles River, Arbresle, France) were randomized, and housed under pathogen-free conditions with standard mice pellet diet and water provided *ad libitum*. After acclimatization for 7 days, mice were injected intramuscularly with K7M2 cells (10^6^ cells/15 µL PBS) in both thighs under isoflurane/air inhalational anesthesia. Muscle infiltrated with tumor tissues and lungs were collected at one month after cell injection, fixed in 4% paraformaldehyde in PBS, and then embedded in paraffin.

### 2.15. Immunohistochemistry

Formalin-fixed paraffin-embedded (FFPE) sections (4 µm thick) dewaxed in xylene were rehydrated through a graded series of ethanol, then stained with haematoxilin-eosin-safran (HES) or processed for IHC staining using a Bond Leica automated immuno-stainer instrument. For the detection of CtBP2 and CYR61, tissue sections were processed for heat-induced antigen retrieval in citrate buffer pH6 and EDTA buffer pH9, respectively. Slides were then incubated for 60 min at room temperature with rabbit anti-CtBP2 antibody (Abcam #ab128871; diluted 1:750) or rabbit anti-CYR61 antibody (Bio-Techne SAS #NB100-356; diluted 1: 500). Slides were then processed with the Bond Polymer Refine Detection kit (#DS9800; Leica Biosystems) or the Rabbit HRP PowerVision Kit (ImmunoVisionTechnologies). The signal was revealed with DAB and counterstained with haematoxylin. For the immunohistochemical detection of hypoxia, tissue sections were incubated for 60 min at room temperature with a rabbit polyclonal anti-GLUT-1 antibody (Clinisciences #RP128; diluted 1:200). The cytoplasmic signal was revealed with the Rabbit HRP PowerVision Kit (ImmunoVisionTechnologies) and with DAB and counterstained with hematoxylin. Whole tumor tissue sections from each sample were digitized using a slide scanner.

### 2.16. Protein-protein interaction network

The STRING database that integrates all known and predicted associations between proteins, including both physical interactions as well as functional associations was used as Core Data Resource (https://string-db.org/; version 11.5). Settings included full STRING network, excluding gene fusion from the selection of active interaction sources.

### 2.17. Statistical analysis

GraphPad Prism Software (version 9.5.1 for Windows; San Diego, CA, USA) was used to determine statistical significance between groups. *P* values <0.05 were considered significant.

## 3. Results

### 3.1. CYR61 expression level influences CtBP2 gene expression

To investigate the modulation of gene expression dependent on CYR61 levels, we compared the transcriptome of K7M2 cells stably overexpressing or silenced for CYR61 to the control cell line. Using selection criteria of adjusted p-value <0.05 *vs.* control cells and |(Fold Change)|>10% cut-off, we identified differentially expressed genes (DEGs) in CYR61-silenced cells and CYR61-overexpressing cells. Among the differentially expressed genes in common in both comparisons only one gene named CtBP2 (C-terminal Binding Protein 2; Gene ID: 1488) presented an opposite variation: downregulation in CYR61-silenced cells (-12%; p=0.039) and upregulation in CYR61-overexpressing cells (+16%, p=0.0128), compared to control cells. Quantitative RT-PCR analyses confirmed the decrease in CtBP2 mRNA levels in CYR61-silenced K7M2 cells (-27%; p=0.028; Figure 1A) and the increase in CYR61-overexpressing K7M2 cells (+2.2-fold; p=0.002). Western blot analyses confirmed the modulations also at a protein level (Figure 1B).

**Figure 1.**
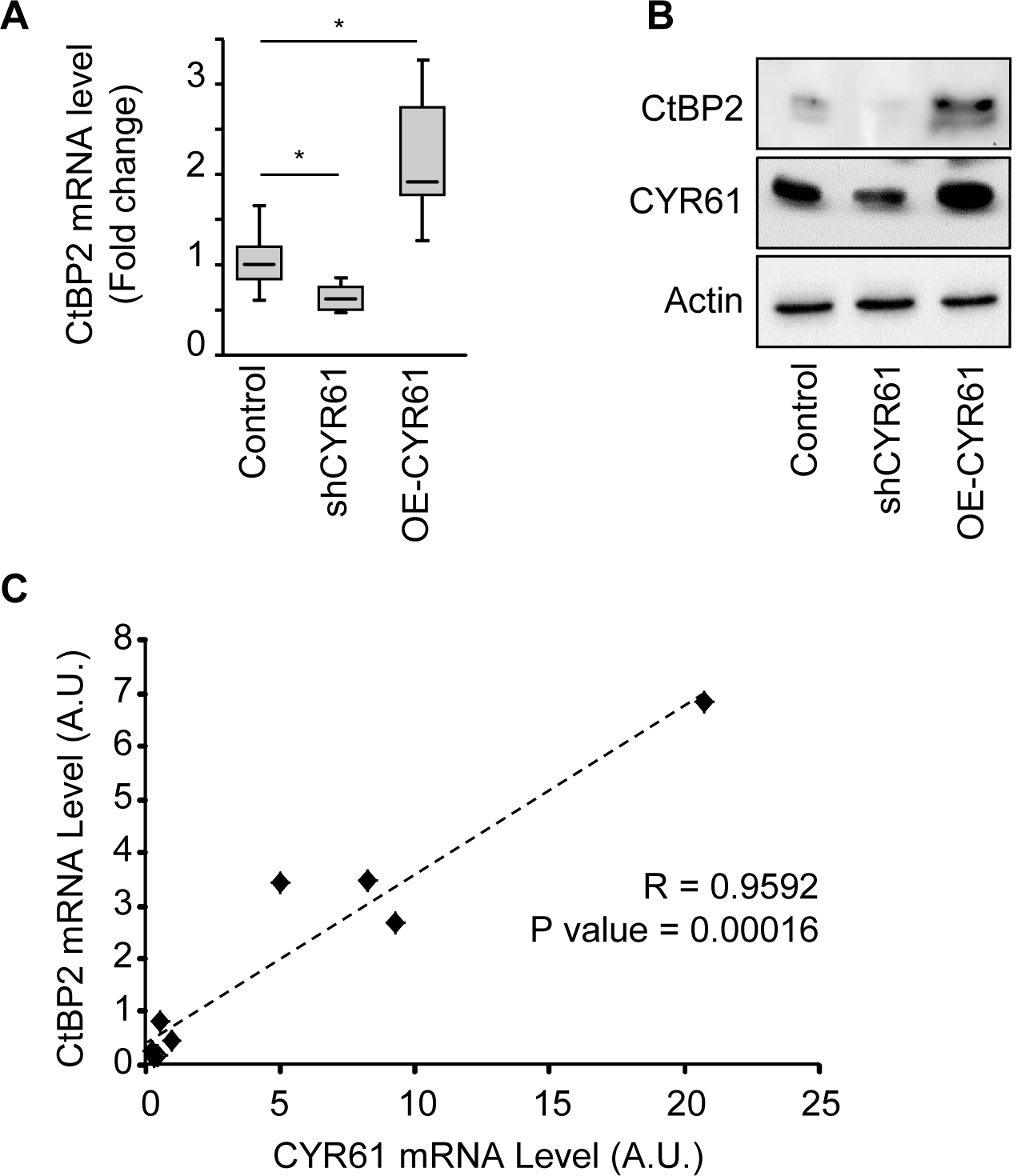
CtBP2 is a transcriptional target of CYR61. **A** Expression pattern of CtBP2 mRNA in K7M2 cell lines modified for CYR61, as assessed by RT-qPCR. GAPDH was used as internal reference gene. The relative mRNA level was calculated using the 2^−ΔΔCT^ method and expressed as box plot (n=3). An asterisk (*) indicates a statistically significant difference (P-value < 0.05). **B** Expression pattern of CtBP2 and CYR61 protein in K7M2 cell lines modified for CYR61, as assessed by western blot. Actin was used as loading control. **C** Spearman correlation between the CtBP2 and CYR61 mRNA level, evaluated by RT-qPCR in a panel of eight human osteosarcoma cell lines. The black dotted line shows the regression line.

We next investigated the correlation between CYR61 and CtBP2 expression levels in a panel of eight human osteosarcoma cell lines. A strong positive correlation was detected between CtBP2 and Cyr61 mRNA levels as assessed by RT-qPCR (Figure 1C).

These results suggest that the expression level of the co-transcription factor CtBP2 correlates to CYR61 expression level in the osteosarcoma cells.

### 3.2. CtBP2 and CYR61 network harbors core stem factors

We next performed *in silico* analysis of CtBP2 and CYR61 potential protein interaction network using the STRING database (https://string-db.org/). Despite the absence of direct physical or functional interaction between CtBP2 and Cyr61, an indirect link through the transcription factors Sox2 (Sex determining region Y-Box 2), Zeb1 (Zinc finger E-Box binding homeobox 1), POU5F1 (POU Class 5 Homeobox 1), and Nanog Homeobox was suggested (Figure 2A and Table 1). We thus evaluated the mRNA levels of the different core stem genes in our panel of eight osteosarcoma cell lines. As expected, the expression levels of Nanog, POU5F1 and Zeb1 strongly positively correlated to each other (Table 2). We also confirmed strong positive correlation between these genes and CtBP2 or Cyr61 expression levels (R>0.8).

**Figure 2.**
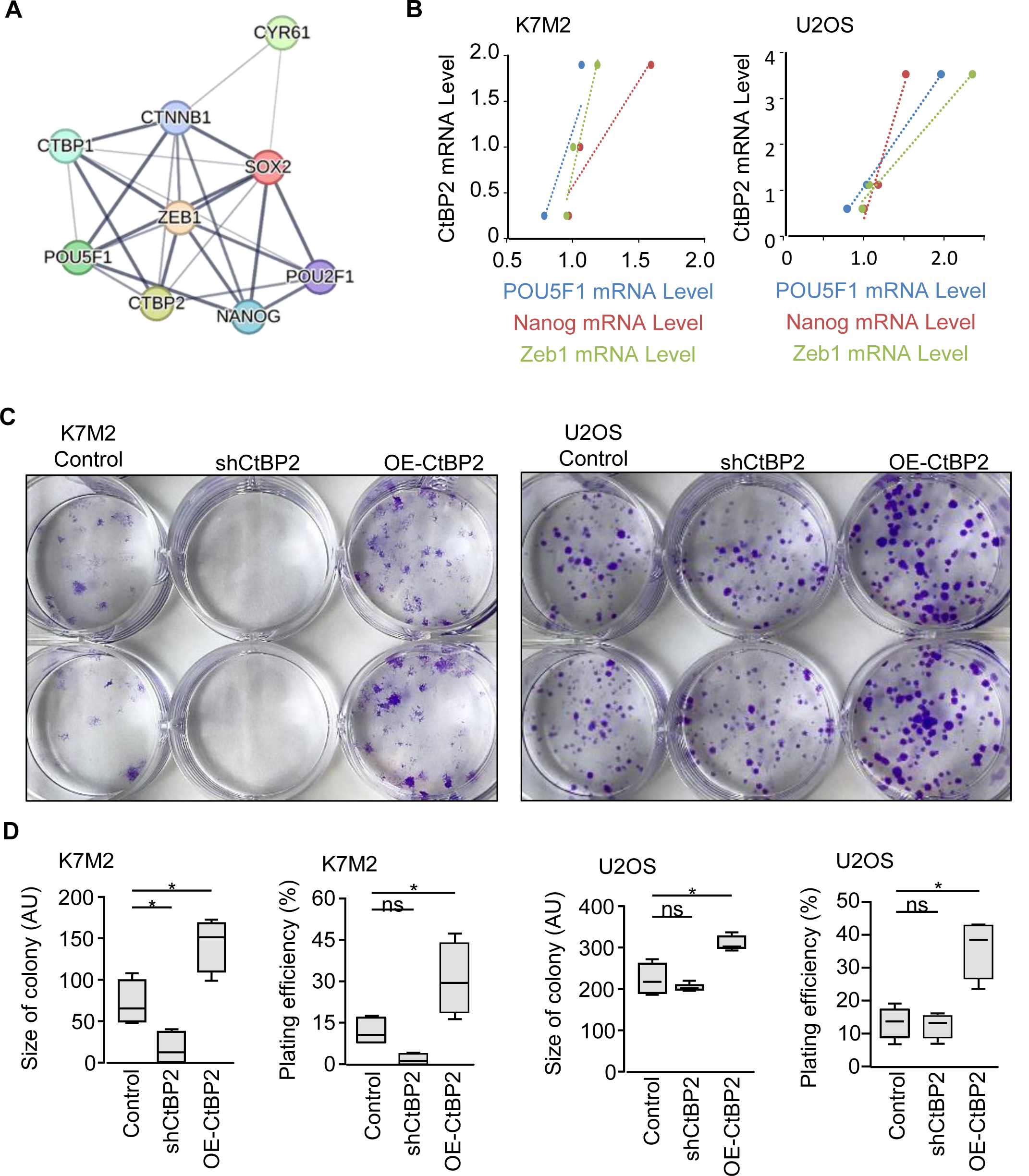
CtBP2 expression level is related to stemness. **A** Interaction network diagram of CtBP2 and CYR61 proteins. Network nodes represent proteins; the edges indicate both functional and physical protein associations; line thickness indicates the strength of data support: the thicker the gray connecting line, the stronger is the predicted interaction between the two proteins. **B** Spearman correlation between the indicated mRNA levels in K7M2 and U2OS modified cell lines, evaluated by RT-qPCR. GAPDH was used as internal reference gene. The relative mRNA level was calculated using the 2^−ΔΔCT^ method and expressed as box plot (n=3 independent experiments). The dotted lines show the regression lines. **C** Representative images from clonogenic assays after staining with crystal violet solution. K7M2 and U2OS cell lines were plated at 100 cells/cm², and incubated for 10 days. **D** Average size and number of colonies formed. Results are expressed as box plot (n=4). An asterisk (*) indicates a statistically significant difference (P-value < 0.05 *vs.* Control).

**Table 1.** Interaction scores from the STRING Database as indicators of confidence in the functional or physical protein interaction. All scores rank from 0 to 1, with 1 being the highest possible confidence.

|  | CYR61 | CtBP2 | SOX2 | ZEB1 | POU5F1 | NANOG |
| --- | --- | --- | --- | --- | --- | --- |
| CYR61 | 1.000 |  |  |  |  |  |
| CtBP2 | - | 1.000 |  |  |  |  |
| SOX2 | 0.403 | 0.537 | 1.000 |  |  |  |
| ZEB1 | - | 0.985 | 0.975 | 1.000 |  |  |
| POU5F1 | - | 0.700 | 0.999 | 0.679 | 1.000 |  |
| NANOG | - | - | 0.998 | 0.960 | 0.999 | 1.000 |

**Table 2.** Matrix correlation of indicated mRNA levels, as assessed by RT-qPCR in a panel of eight human osteosarcoma cell lines.

|  | CYR61 | CtBP2 | SOX2 | ZEB1 | POU5F1 | NANOG |
| --- | --- | --- | --- | --- | --- | --- |
| CYR61 | 1.000 |  |  |  |  |  |
| CtBP2 | 0.917 | 1.000 |  |  |  |  |
| SOX2 | 0.967 | 0.829 | 1.000 |  |  |  |
| ZEB1 | 0.959 | 0.847 | 0.934 | 1.000 |  |  |
| POU5F1 | 0.975 | 0.889 | 0.978 | 0.967 | 1.000 |  |
| NANOG | 0.995 | 0.896 | 0.962 | 0.981 | 0.976 | 1.000 |

These results suggest that Cyr61 and CtBP2 expression correlates with the expression of stemness markers.

### 3.3. CtBP2 expression level influences osteosarcoma cell stemness

To investigate the role of CtBP2 in osteosarcoma cells, we established new cell lines either repressing or over-expressing CtBP2 by cell transduction with lentiviral vectors. As expected, the stable genomic integration of CtBP2 targeting shRNA sequences led to a reduction in CtBP2 mRNA levels, as compared to cells transduced with non-relevant shRNA sequences (-63%, p=0.002 for K7M2 cell line and -82%, p<0.001 for U2OS cell line; Supplemental Figure 1A). Likewise, the stable genomic integration of the full length CtBP2 coding sequence led to an increased CtBP2 mRNA levels, as compared to control cells (2.5-fold for K7M2 cell line, p=0.017 and 3.5-fold for U2OS cell lines, p=0.006). Western blot analyses confirmed the stable modulation of CtBP2 expression at a protein level (Supplemental Figure 1B).

We first evaluated the expression levels of stem cell markers in the CtBP2-modified cell lines. As observed for the panel of cell lines, the mRNA levels of Zeb1, POU5F1 and Nanog positively correlated to those of CtBP2 (R>0.8; Figure 2B). We next performed a clonogenic assay. Interestingly, the ability of individual cells to form colony was clearly dependent on CtBP2 expression levels. Indeed, the repression of CtBP2 correlated with reduced number and size of colonies, whereas the CtBP2-overexpression promoted the colony formation ability (Figure 2C-D). We then investigated the influence of CtBP2 levels on *in vitro* cell characteristics. Cell metabolic activity assessed by the MTT test (supplemental Figure 1C), DNA replication rate assessed by the BrdU incorporation assay (supplemental Figure 1D), and cell morphology characterized by measurement of cell spreading area, perimeter, and shape/circularity (supplemental Figures 1E-F) were stable among the modified cell lines.

These results suggest that variations in CtBP2 expression levels influence stemness markers levels in osteosarcoma cells, but does not cause changes in cell morphology, viability or proliferation.

### 3.4. CtBP2 controls *in vitro* osteosarcoma cell migration

We compared the cell migration capacity of CtBP2-silenced and -overexpressing cells using modified Boyden chamber assay. After 10 h of incubation a total of 27 ± 10 % of shCtBP2 cells, and 67 ± 21 % of CtBP2 overexpressing cells migrated through the membrane (p=0.0016; Figures 3A-B). This ratio of more than twice as many migrating OE-CtBP2 cells than shCtBP2 cells was confirmed using the modified U2OS human cell lines (p<0.001; supplemental Figures 2A-B). A wound healing assay confirmed the CtBP2 levels-dependent cell motility. Indeed, the number of cells migrating into the cell-free scratch region was reduced in CtBP2-silenced population (-35%, p<0.001; Figures 3C-D), and increased in CtBP2-overexpressing population (+23%, p<0.001), leading to an average OE/sh ratio of 1.9-fold for K7M2 cell lines. Again, the U2OS modified cell lines presented similar behavior: -19% (p=0.0018) for CtBP2-silenced cells and +27% (p=0.0048) for CtBP2-overexpressing cells, leading to an average OE/sh ratio of 1.57-fold (Supplemental Figures 2C-D).

**Figure 3.**
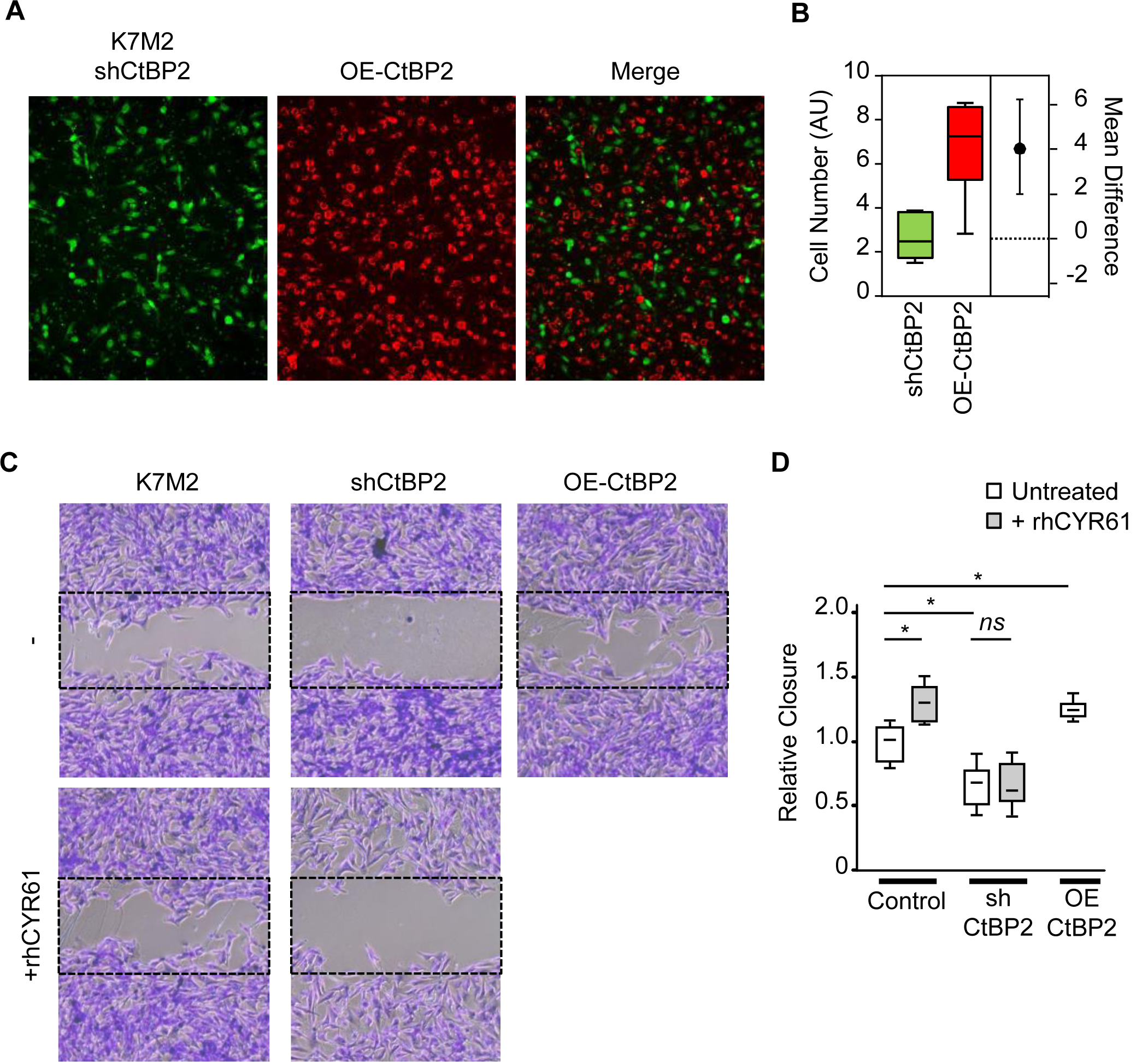
CtBP2 favors cell migration. **A** Representative images of K7M2 cell migration after 6 h incubation, as assessed by Boyden chamber assay. CtBP2-silenced K7M2 cells were stained in green, and CtBP2-overexpressing cells were stained in red before seeding. **B** Estimation plot representing the relative number of migrating cells. The difference between the group means is represented on the right. **C** Representative images of cell migration (wound healing assay) taken at time 18 h after the wound. Control and CtBP2-silenced K7M2 cells were incubated in the presence or absence of recombinant CYR61 (1 µg/ml). The dotted rectangles outline the initial wound surface. **D** Quantitative evaluation of the cell migration rate. Results are expressed as box plot (n=6). An asterisk (*) indicates a statistically significant difference (P-value < 0.05 *vs.* control).

We next investigated the involvement of CtBP2 in the CYR61-dependent cell migration process we previously described (*6, 16, 17, 23*). As expected, the supplementation of culture medium with recombinant CYR61 increased the number of migrating K7M2 cells (+30%, p=0.0252 *vs.* untreated; Figures 3C-D). Interestingly, recombinant CYR61 supplementation did not significantly modify CtBP2-silenced cell migration rate (p=0.4617). Correspondingly, supplementation with recombinant CYR61 increased the number of migrating U2OS cells (+28%, p=0.0026; supplemental Figures 2C-D), but not of CtBP2-silenced cells (p=0.1890).

Taken together, these results suggest that CtBP2 upregulation supports osteosarcoma cell migration. Moreover, these results indicate that osteosarcoma cell motility requires minimal CtBP2 expression levels and that CYR61-mediated cell migration is at least in part dependent on CtBP2 upregulation.

### 3.5. CtBP2 controls *in vitro* osteosarcoma cell invasiveness

We investigated the influence of CtBP2 expression levels on cell invasiveness. We evaluated the MMP2 activity that plays an important role in the invasive process of osteosarcoma cells (*3, 4, 6, 17*). CtBP2-silenced K7M2 cells exhibited a reduced MMP2 activity compared to control cells (-17%, p=0.0012; Figure 4A), whereas CtBP2-overexpressing cells exhibited a higher MMP2 activity (+27%, p=0.0083). In the same way, CtBP2-silenced U2OS cells exhibited a reduced MMP2 activity compared to control cells (-24%, p=0.0016), whereas CtBP2-overexpressing cells exhibited higher MMP2 activity (+57%, p<0.0001).

**Figure 4.**
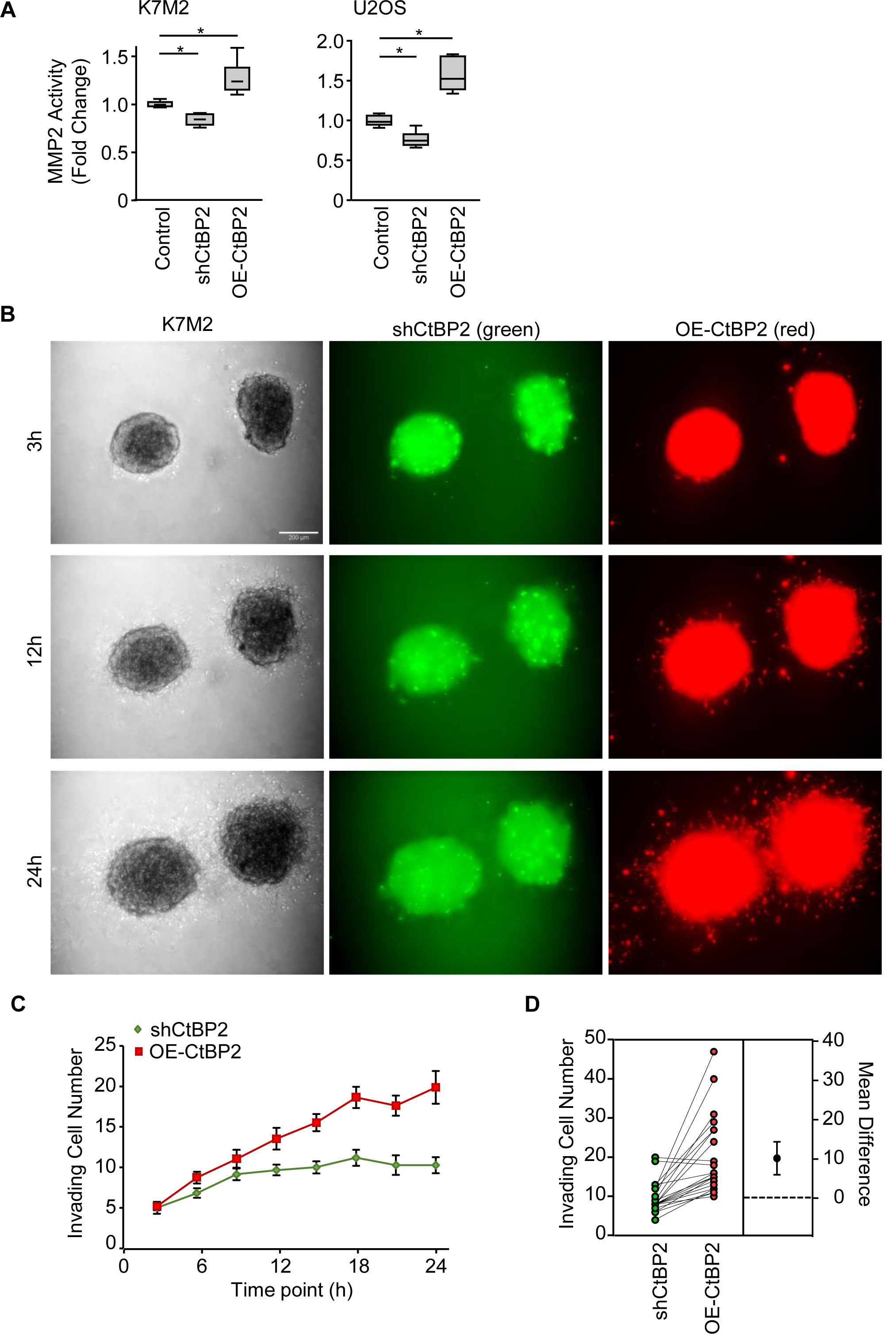
CtBP2 favors cell invasion through extracellular matrix. **A** Relative Matrix Metalloproteinase-2 (MMP2) activity in K7M2 and U2OS modified cell lines, as assessed by colorimetric assay. Results are expressed as box plot (n=10). An asterisk (*) indicates a statistically significant difference (P-value < 0.05 *vs.* Control). **B** Representative images of spatial spheroid invasion assays. 3D cultured spheroid made of mix (1:1) of CtBP2-silenced and CtBP2-overexpressing K7M2 were embedded in Matrigel basement membrane, and incubated for 24 h (bar = 200 µm). CtBP2-silenced K7M2 cells were stained in green, and CtBP2-overexpressing cells were stained in red. **C** Quantitative evaluation of the cells invading the Matrigel basement membrane at different time points. Results are expressed as mean ± standard error of the mean (n=23). **D** Estimation plot representing the relative number of invading cells as compared to Control. The difference between the group means is represented on the right.

We next investigated the ability of the modified cell lines to invade an extracellular matrix. 3D condensed structures (spheroids) established by homogeneously mixing pre-stained CtBP2-silenced and -overexpressing cells (1:1 ratio) were incubated in the presence of Matrigel basement membrane. Cells penetrating into the surrounding matrix environment were monitored and scored at different time points (Figures 4B-C). CtBP2-overexpressing cells that escape the spheroid are more numerous than CtBP2-silenced cells, representing about two third of invading cells after 24 h (p<0.0001; Figure 4D).

These results indicate that CtBP2 overexpression favors Matrix Metalloproteinase activity and osteosarcoma cell invasion.

### 3.6. Overexpression of CtBP2 at the invasive front of the tumors

To confirm the involvement of CtBP2 in tumor dissemination we used syngeneic tumor cells graft models. We implanted K7M2 cells into tight muscle of BALB/c-*nu* mice to establish primary osteosarcoma tumors. IHC staining on FFPE sections of Control cells-derived tumors revealed a heterogeneity in the CtBP2 signal intensity within the tissue section (Figure 5A). Indeed, osteosarcoma cells invading the surrounding normal tissue exhibited a higher signal intensity for CtBP2 as compared to cells located in a more central area (+79%; p=0.0002; Figure 5B). We thereafter investigated CtBP2 signal intensity in FFPE sections of tumor samples derived from the CYR61-modified cells. Surprisingly, no significant difference was detectable in CtBP2 signal intensity at the central area of tissue sections between the different models (p>0.3). Nevertheless, a similar profile to the Control group was detected for the CYR61-overexpressing model, with an increase in CtBP2 signal intensity at the invasive front as compared to the central area (+84%; p=0.0002; Figures 5A-B, Supplemental Figure 3). In contrast, CtBP2 signal in cells located at the front area of the tumor derived from CYR61-silenced grafted cells was weakly increased as compared to the central area (+29%; p=0.0178).

**Figure 5.**
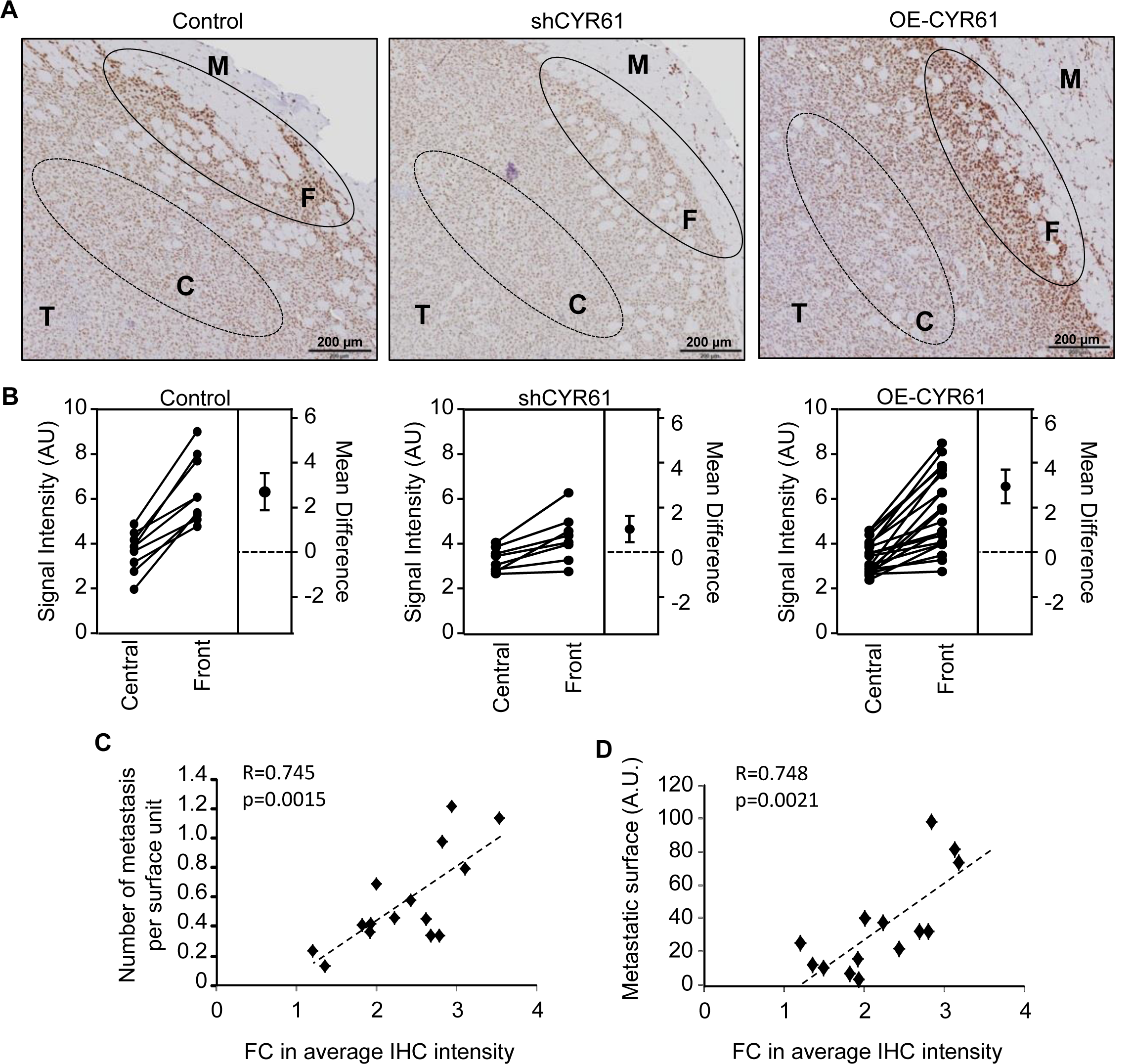
CtBP2 induction by CYR61 is maximal at the invasive front of the tumor. **A** CtBP2 immunohistochemical staining of FFPE tissue sections of Control, CYR61-silenced or CYR61-overexpressing cell line derived xenografts. Tumor (T) and surrounding muscle (M) as well as the tumor core area (C) and invasive front (F) are indicated. The scale bars represent 200 µm. **B** Estimation plots representing the relative CtBP2 signal intensity determined for Control, CYR61-silenced or CYR61-overexpressing cell line derived xenografts. Results are expressed as paired sample comparison between the tumor core and invasive front. The difference between the means are represented on the right of each plot. **C** Spearman correlation between the average CtBP2 signal intensity and the number of metastatic foci in the lungs. The dotted line shows the regression line. **D** Spearman correlation between the average CtBP2 signal intensity and the lung metastatic surface. The dotted line shows the regression line.

We also evaluated the relationship between the variation in CtBP2 expression levels and the occurrence of metastases. A strong positive correlation was detected between CtBP2 upregulation at the invasive tumor front, and the number or surface of pulmonary metastatic foci (Figures 5C-D).

These results suggest an induction of CtBP2 expression in CYR61-expressing osteosarcoma cells at the invasive front that correlates with the metastatic dissemination process. Moreover, there is a distinct blockade of the CYR61-dependent induction of CtBP2 expression in more central region of the tumors.

### 3.7. Micro-environmental conditions impair CtBP2 induction

To confirm the importance of the spatial location of cells into the tumor mass for the link between CYR61 and CtBP2 expression levels, we established 3D bigger spheroids. As suspected, we observed a gradient for CtBP2 signal intensity with a higher level in the cells located at the outer proliferating envelope of the spheroid FFPE sections as compared to the necrotic core (average +10%, p=0.0220; Figures 6A-B). We evaluated the expression levels of GLUT1 as marker to detect hypoxia. We observed a gradient with a lower expression level in the cells located at the outer layer of the spheroid as compared to those in the center (average -8%, p=0.0288). Globally, we detected a strong negative correlation between CtBP2 and GLUT1 signal intensities (Figure 6C). To test how oxygen levels influence CtBP2 expression, we incubated K7M2 cells as 2D monolayers under either normoxic or hypoxic conditions (21% or 3% oxygen, respectively). As expected, we observed a marked decrease in CtBP2 expression levels in cells cultured under hypoxia as compared to normoxia at both mRNA and protein levels (Figure 6D-E).

**Figure 6.**
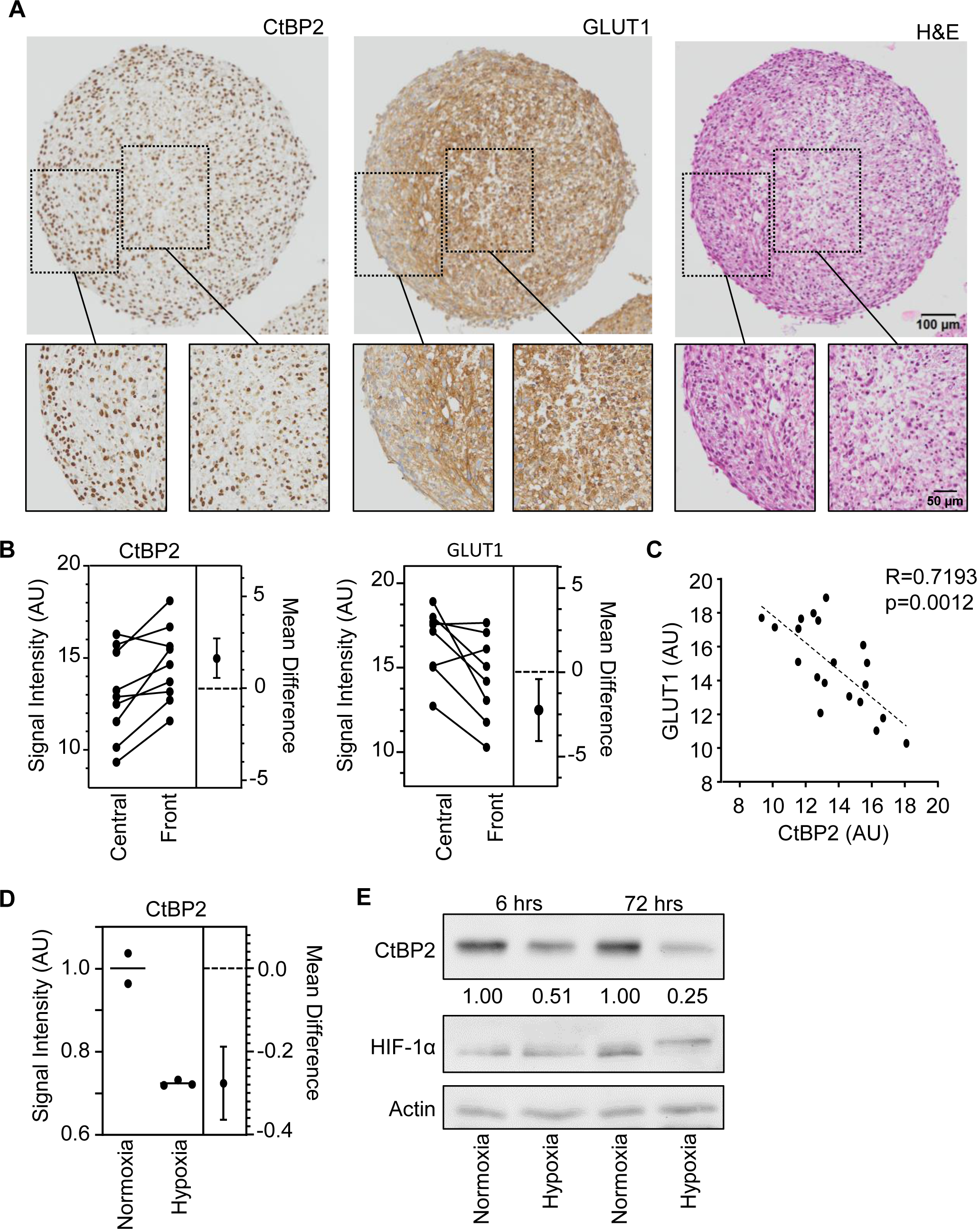
CtBP2 expression is dependent on oxygen levels. **A** Immunohistochemical staining for CtBP2 and GLUT1, and H&E staining of FFPE section of K7M2 cultured as 3D spheroid. The scale bars represent 100 µm or 50 µm for below insert. **B** Estimation plots representing the relative CtBP2 and GLUT1 signal intensity. Results are expressed as paired sample comparison between the spheroid central and front area. The difference between the means are represented on the right of each plot. **C** Spearman correlation between CtBP2 and GLUT1 signal intensities. The dotted line shows the regression line. **D** Expression pattern of CtBP2 and HIF-1α protein in K7M2 cells cultured as 2D monolayer for 6 h or 72 h in normoxic or hypoxic condition, as assessed by western blot. Actin was used as loading control.

These results suggest that hypoxic conditions interfere with CtBP2 induction of expression, and consequently that non-hypoxic conditions such as external location of tumor cells aggregates favor it.

## 4. Discussion

A major negative prognostic factor for survival of osteosarcoma patients is the presence of metastasis at diagnosis (*24*). Preventing the metastatic dissemination is thus a crucial issue for this pediatric cancer. Us and others have previously demonstrated that CYR61 is a key pro-metastatic factor in bone tumor cells (*6, 16, 17*). While no clear difference was detectable in the primary tumor growth rate, the metastatic dissemination rate positively correlates with CYR61 expression levels. CYR61 is secreted, extracellular matrix (ECM)-associated signaling protein, reported to solicit several key membrane receptors such as TGFβR, VEGFR, integrin αvβ3 or IGF1Rβ (*reviewed in* (*25*)). The resulting signaling cascades involve various intracellular effectors such as PI3K-Akt / ERK1/2-MAPKs / NFkB / ILK / TEAD-YAP, and lead to the transcriptional activation of genes related to proliferation, migration, invasion and angiogenesis.

The present study identified for the first time CtBP2 (C-terminal Binding Protein 2) as a transcriptional target of CYR61. The CtBP family of proteins are transcriptional co-regulators implicated in mammalian embryogenesis and development, and oncogenesis (*reviewed in* (*26*)). The resulting regulative activity relies on their interacting partners (*27*). Thus, CtBPs exhibit a predominant co-repressor activity thereby contributing to the negative regulation of the expression of many tumor suppressor genes (*reviewed in* (*28*)). However, CtBPs also exhibit co-activating activity favoring the expression of genes that promote proliferation and cancer stem cell self-renewal, or the epithelial-mesenchymal transition (EMT) (*29*). We previously reported the induction of an EMT-like process in osteosarcoma cells overexpressing CYR61 (*17*). *In silico* analysis suggested that the CYR61 and CtBP2 interaction network may involve stemness markers, namely Sox2, Nanog, Zeb1 and POU5F1. We confirmed a strong positive correlation (R > 0.8) between CYR61, CtBP2 and the stemness markers expression at the transcriptional levels in our osteosarcoma cells. Moreover, our functional validation experiments suggest that CtBP2 favors the acquisition of stem cell-like characteristics. These results reinforce our hypothesis that CtBP2’s input is essential in the CYR61-dependent induction of EMT-like process of osteosarcoma cells, and is required for tumor cell dissemination. To our knowledge, no available data foresees a link between CtBP2 and CYR61 in any type of tumor or normal cells.

According to the involvement of CtBP2 in osteosarcoma cells behavior, only one publication recently reported a control of the expression of genes related to DNA repair by a transcriptional complex comprising CtBP1/2 heterodimer, CtBP-interacting protein (CtIP), histone deacetylase 1 (HDAC1), and two subunits of activating protein 1 (AP1) (*30*). The genomic chaos occurring in osteosarcoma cells drives the heterogeneity and complexity of this cancer. The random but extensive DNA rearrangements, named *chromothripsis*, (*31*), are a crucial issue halting the development of therapies as complement to chemotherapy that are frequently inefficient.

The metastatic spread to distant organs is a second key concern. Here, we report a positive correlation between CtBP2 levels in osteosarcoma cells and their *in vitro* motility and *in vivo* invasiveness capacities. We also provide a proof of concept that CtBP2-silencing lowered 2D and 3D *in vitro* cell migration capacity, and prevented CYR61-induction of cell migration. Moreover, we reported that CtBP2-silencing significantly impairs *in vivo* tumor dissemination capacity.

Overall, we revealed CtBP2 as a downstream transcriptional target of CYR61, and demonstrated that CtBP2 expression is required for CYR61-dependent pro-metastatic dissemination of osteosarcoma, by favoring cell migration and invasiveness. In a very interesting manner, we observed that CtBP2 repression (silencing) discontinued the pro-metastatic cascade initiated by CYR61. CtBP2 thus represents an interesting potential target for the prevention of tumor spreading. This factor is not restricted to osteosarcoma as hyperactivity of CtBPs mediates oncogenic functions and promotes aggressive and metastatic neoplastic behavior in various solid tumors (reviewed in (*32*)). In that way, compounds interfering with either protein-protein interaction or dehydrogenase enzymatic activity were tested with limited success in *in vivo* assays. The development of derivative or related molecules is on the way, and such inhibitors could represent a valuable option for osteosarcoma.

Another valuable point that we highlighted in our osteosarcoma cells is that the tumor cell location within the tumor mass is critical for the good execution of the CYR61-CtBP2-migration/invasion cascade. Indeed, the highest levels of CtBP2 expression were detected in the surrounding cells whereas no upregulation of CtBP2 was detectable in the cells located in a more central area. The presence of hypoxia responsive elements (HRE) in the CtBP2 promoter (*33*) suggests that HIF family members interfere with the induction of CtBP2 expression. In fact, CtBP has been described as critical mediator of the oxygen-sensing response (*34, 35*). Our *in vitro* experiments confirmed the role of micro-environmental conditions such as the oxygen levels in the regulation of CYR61-dependent induction of CtBP2 expression by osteosarcoma cells.

## 5. Conclusions

Clinical management of osteosarcoma requires new therapeutic strategies to prevent the development of metastatic disease, and thus improve the outcomes for patients with poor prognosis. Our previous studies as well as that of others strongly suggest CYR61/CCN1 as a prognosis biomarker of metastatic dissemination. The present study provides new interesting information on the CYR61-driven pro-metastatic cascade, involving the transcriptional induction of the co-repressor CtBP2. Our data also revealed that this cascade occurs under oxygen-driven conditions, as encountered rather at a peripheral location of the tumor, representing the invasive front. CtBP2 is indeed required for pro-metastatic dissemination of osteosarcoma, by favoring cell migration and invasiveness. Hence, CtBP2 is a crucial element of the CYR61-driven invasiveness of osteosarcoma cells.

## Abbreviations

BrdU: Bromo-deoxyuridine
CtBP2: C-terminal Binding Protein-2
CTGF: Connective Tissue Growth Factor
CYR61: Cysteine Rich Protein-61
EMT: Epithelial to Mesenchymal Transition
FFPE: Formaldehyde Fixed Paraffin Embedded
GAPDH: Glyceraldehyde-3-Phosphate Dehydrogenase
GLUT1: Glucose Transporter 1
H&E: Hematoxylin and Eosin
HIF-1α: Hypoxia Inducible Factor 1 Subunit Alpha
IHC: Immuno-Histochemistry
MMP: Matrix Metalloproteinase
MTT: 3-(4,5-dimethylthiazol-2-yl)-2,5-diphenyltetrazolium bromide
NANOG: Homeobox Transcription Factor Nanog
NOV: Nephroblastoma Overexpressed
OE-CtBP2: overexpression of CtBP2
OE-CYR61: overexpression of CYR61
pHEMA: Poly(2-hydroxyethyl methacrylate)
POU5F1: POU class 5 Homeobox 1
sh-CtBP2: silencing of CtBP2
sh-Cyr61: silencing of Cyr61
SOX2: SRY (Sex determining Region Y)-Box 2
WISP: WNT1-Inducible-Signaling Pathway
ZEB1: Zinc finger E-Box binding homeobox 1

## Acknowledgments

We thank the Pre-clinical Evalution Platform for their technical assistance with the animal work (UMS AMMICa, Gustave Roussy, Villejuif, France). We thank Dr Nicolas Signole, Olivia Bawa and Hélène Rocheteau (Experimental Pathology Platform of Gustave Roussy), and Dr Emilie Louvet (present address AstraZeneca Oncology iMED, London, UK) for their help in IHC execution and analyses. We thank Dr Francesco Baschieri (Inserm U1279, Villejuif, France) for its active contribution in 3D models establishment and imaging.

This work was supported by Inserm (France), and Fondation de l’Avenir pour la Recherche Médicale Appliquée grant ET688 (France). Dr L. Di Patria was the recipient of a visiting grant from the University of Urbino Carlo Bo (Italy) to complete her doctorate course.

## Author Contributions

LDP, NH, RO, BS and OF carried out the experiments and analyzed the data. BS and OF conceived and supervised the project, wrote and revised the manuscript. All authors read and approved the final manuscript.

## Ethics approval

Animal experiments and procedures received the approval of the local French ethical animal committee of Paris Saclay University (CEEA 26).

## Supporting Information

**Supplemental Table S1:**
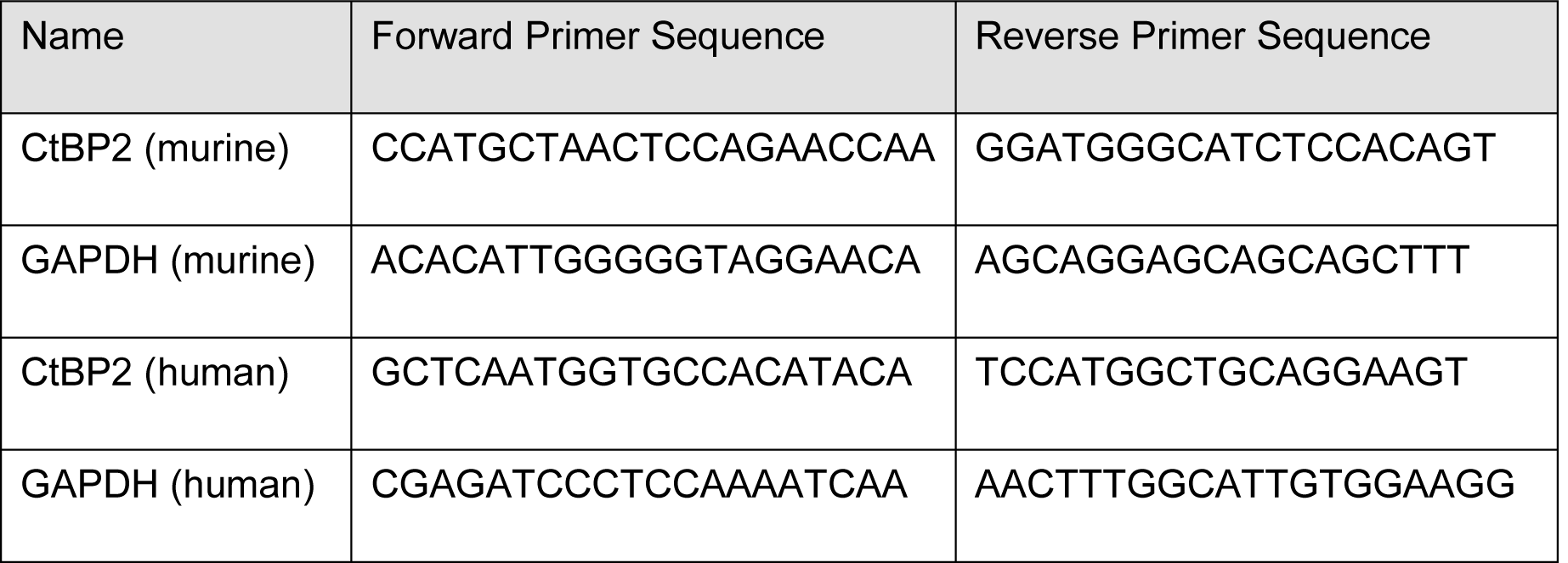
Specific primers sequences for quantitative PCR assays.

**Supplemental Table S2:** Antibodies for western blot and immunohistochemistry staining.

| Name | Reference | Application | Dilution |
| --- | --- | --- | --- |
| CtBP2 | #ab128871 (Abcam) | IHC | 1:750 |
| CtBP2 | #ab128871 (Abcam) | WB | 1:1000 |
| CYR61 | #NB100-356 (Bio-Techne SAS) | IHC | 1:500 |
| CYR61 | #NB100-356 (Bio-Techne SAS) | WB | 1:1000 |
| GLUT1 | #RP128 (Clinisciences) | IHC | 1:200 |
| GAPDH | #MAB374 (Millipore) | WB | 1:1000 |

**Supplemental Figure S1.**
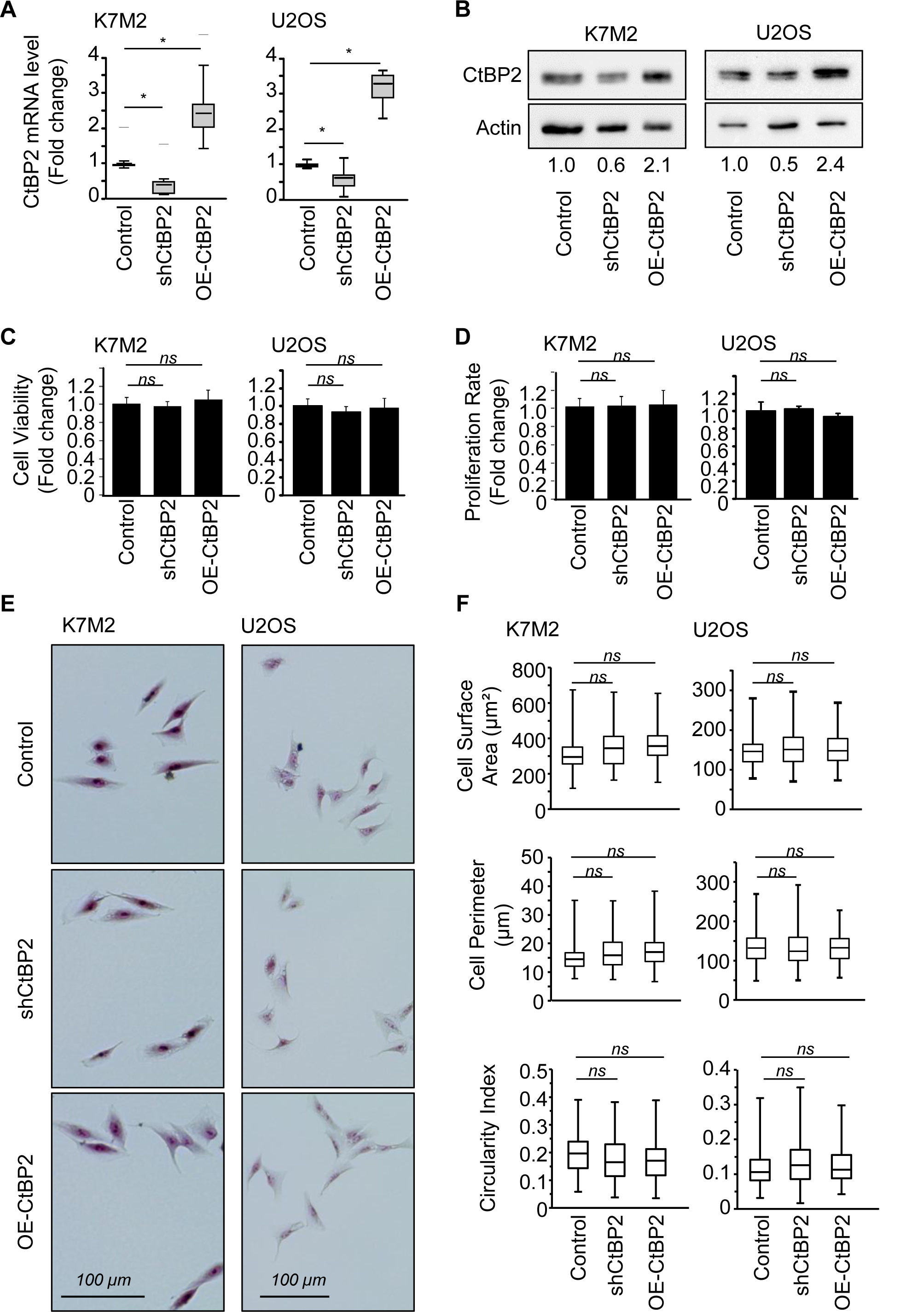
**A** Expression pattern of CtBP2 mRNA in Control and stably modified cell lines, as assessed by RT-qPCR. GAPDH was used as internal reference gene. The relative mRNA level was calculated using the 2^−ΔΔCT^ method and expressed as box plot (n=3 independent experiments). An asterisk (*) indicates a statistically significant difference (P-value < 0.05 *vs.* Control). **B** Expression pattern of CtBP2 protein in K7M2 and U2OS modified cell lines, as assessed by western blot. Actin was used as loading control. **C** Relative cell metabolic activity, as assessed by the MTT assay. Results are expressed as mean ± standard deviation (n=30). **D** Relative DNA replication rate, as assessed by the BrdU incorporation assay. Results are expressed as mean ± standard deviation (n=17). **E** Representative images of Control and stably modified cells. **F** Distribution of cell shape parameters (cell surface, cell perimeter, and circularity index) for the Control and stably modified K7M2 and U2OS cells. Results are expressed as box plot.

**Supplemental Figure S2.**
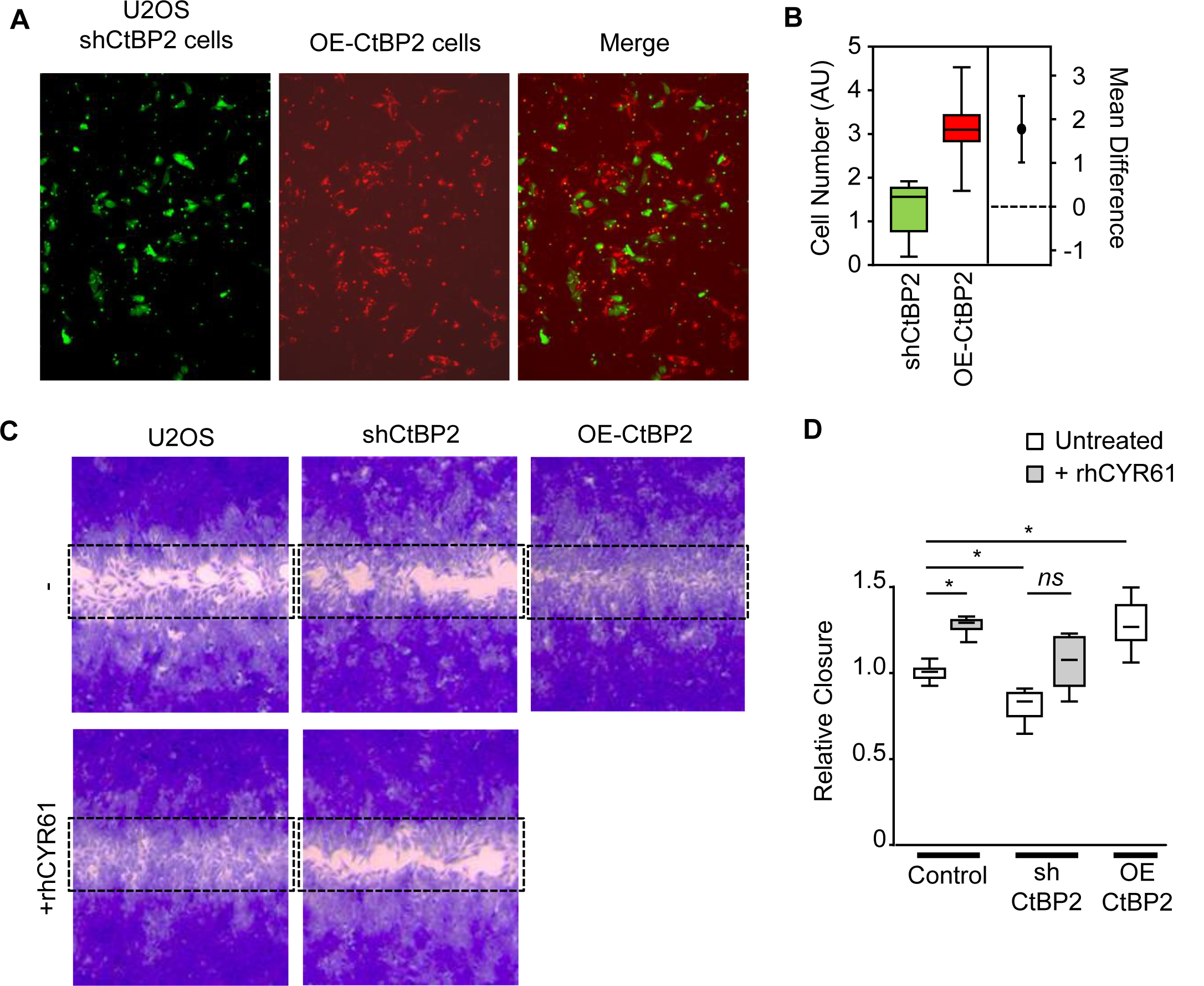
**A** Representative images of U2OS cell migration after 18 h incubation, as assessed by Boyden chamber assay. CtBP2-silenced U2OS cells were stained in green, and CtBP2-overexpressing cells were stained in red. **B** Estimation plot representing the relative number of migrating cells as compared to Control. The difference between the group means is represented on the right. **C** Representative images of cell migration (wound healing assay) taken at time 18 h after the wound. Control and CtBP2-silenced U2OS cells were incubated in the presence or absence of recombinant CYR61 (1 µg/ml). The dotted rectangles outline the initial wound surface. **D** Quantitative evaluation of the cell migration rate. Results are expressed as box plot (n=6). An asterisk (*) indicates a statistically significant difference (P-value < 0.05 *vs.* control).

**Supplemental Figure S3.**
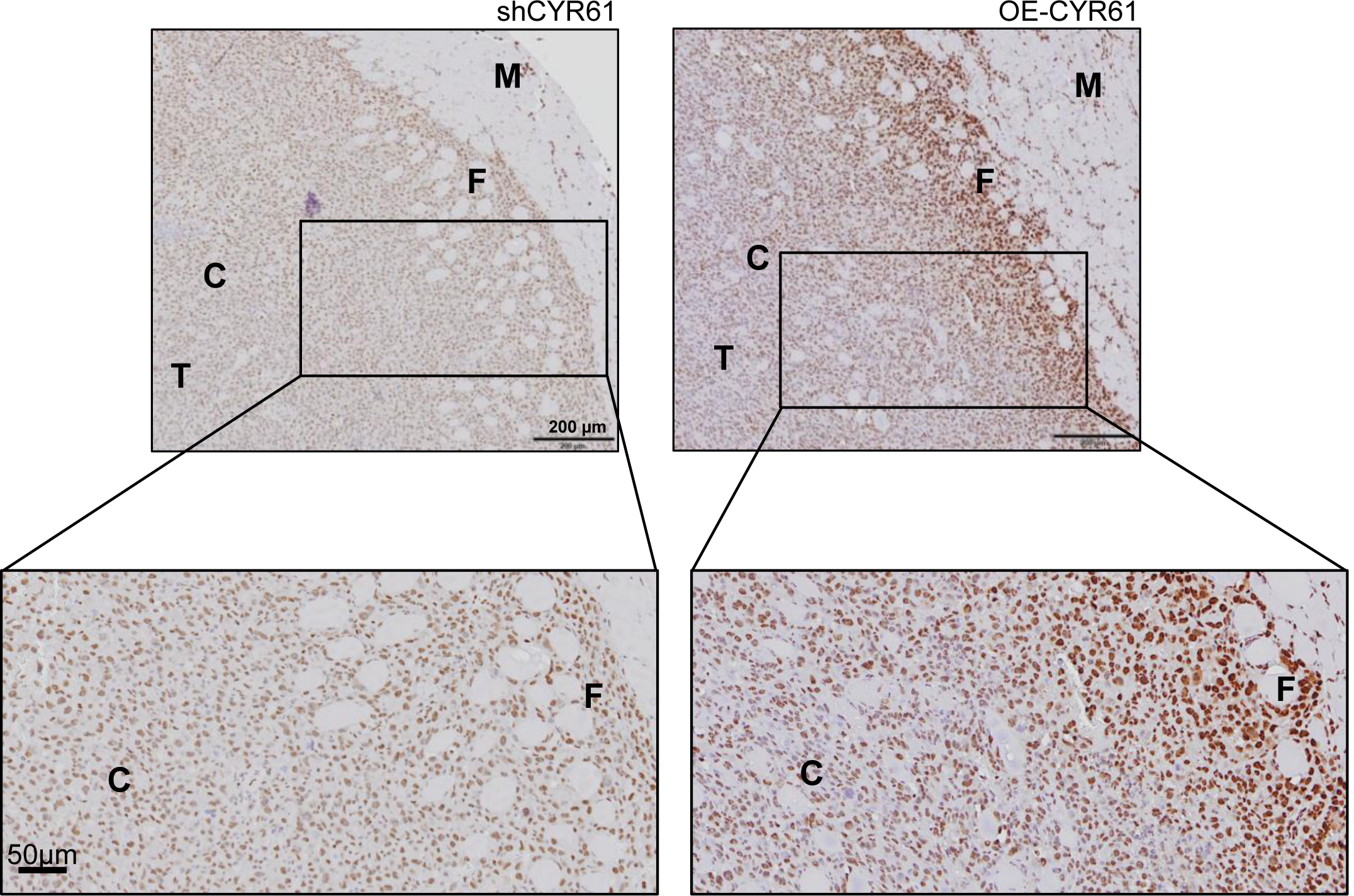
Representative images with higher magnification of CtBP2 immunohistochemical staining of FFPE tissue sections of CYR61-silenced or CYR61-overexpressing cell line derived xenografts. Tumor (T) and surrounding muscle (M) as well as the tumor core area (C) and invasive front (F) are indicated. The scale bars represent 200 µm.

